# Sphingolipid metabolism is spatially regulated in the developing embryo by *SOXE* genes

**DOI:** 10.1101/2023.08.10.552770

**Authors:** Michael L. Piacentino, Aria J. Fasse, Alexis Camacho-Avila, Ilya Grabylnikov, Marianne E. Bronner

## Abstract

During epithelial-to-mesenchymal transition (EMT), significant rearrangements occur in plasma membrane protein and lipid content that are important for membrane function and acquisition of cell motility. To gain insight into how neural crest cells regulate their lipid content at the transcriptional level during EMT, here we identify critical enhancer sequences that regulate the expression of *SMPD3*, a gene responsible for sphingomyelin hydrolysis to produce ceramide, which is necessary for neural crest EMT. We uncovered three enhancer regions within the first intron of the *SMPD3* locus that drive reporter expression in distinct spatial and temporal domains, together collectively recapitulating the expression domains of endogenous *SMPD3* within the ectodermal lineages. We further dissected one enhancer that is specifically active in the migrating neural crest. By mutating putative transcriptional input sites or knocking down upstream regulators, we find that the SoxE-family transcription factors Sox9 and Sox10 regulate the expression of *SMPD3* in migrating neural crest cells. Together these results shed light on how core components of developmental gene regulatory networks interact with metabolic effector genes to control changes in membrane lipid content.

**Highlights:**

- *SMPD3* is expressed in the neural tube, neural crest, and notochord during early development
- *SMPD3* expression is regulated by at least three intronic enhancers
- Sox10 and its binding sites are required for expression by a migratory neural crest-specific *SMPD3* enhancer
- Sox10 is a positive regulator of endogenous *SMPD3* expression during neural crest migration

## Introduction

As cells in the developing embryo transition between behavioral states, such as during an epithelial-to-mesenchymal transition (EMT) and gain of motility, they coordinate significant rearrangements in their plasma membrane protein and lipid content. These changes are necessary for productive cell signaling, new adhesions, and the formation of membrane protrusions. Despite the fact that lipid content plays an important role in membrane function, we lack a comprehensive understanding of how cells in the developing embryo regulate their lipid content at the transcriptional level.

Cells produce a vast array of lipid species used for energy storage, but also for structural components of cellular membranes, and as bioactive signaling molecules. Of these lipid species, the sphingolipids, including sphingomyelin and ceramide, represent a significant fraction of the eukaryotic lipidome (Levental et al., 2017; Lorent et al., 2020; Meng et al., 2021; Sampaio et al., 2011). Sphingomyelin and ceramide production regulate cell growth, survival, and apoptosis, as well as immunological and neurological signaling (Hannun and Obeid, 2018; Shamseddine et al., 2015; Taniguchi and Okazaki, 2020; Wu et al., 2010). More recently, roles for sphingomyelin and ceramide have extended to invasive behaviors during cell migration and cancer metastasis (Canals et al., 2020; Edmond et al., 2015; Levade et al., 2015; Liang et al., 2023). Since these lipids are semi-ubiquitous components of living cells, many enzymes involved in their synthesis and modification are often broadly expressed and regulated post-translationally through transient subcellular localization strategies and conformational changes (Airola et al., 2017; Liang et al., 2023; Shamseddine et al., 2015; Tani and Hannun, 2007; Tomiuk et al., 2000; Wu et al., 2010). Other lipid pathway components, however, exhibit more tissue-specific expression.

One example of a developmentally expressed gene involved in lipid metabolism is *SMPD3*, which encodes a sphingomyelin-hydrolyzing enzyme that we recently showed is expressed specifically within the notochord, neural tube, and neural crest, but absent from the adjacent mesoderm and presumptive epidermis, in the developing chicken embryo (Piacentino et al., 2022). Although neural crest cells originate within the embryonic ectoderm, they subsequently undergo an EMT to delaminate and migrate throughout the periphery before differentiating into diverse cell types (Bronner and Simões-Costa, 2016; Martik and Bronner, 2017; Piacentino et al., 2020; Rogers and Nie, 2018). Early in development, *SMPD3*-dependent ceramide production is critical for neural crest EMT and invasion (Piacentino et al., 2022). This raises the intriguing question of how lipid production may be differentially regulated during development to facilitate EMT and cell migration, and subsequent morphogenesis.

While we have a deep understanding of the gene regulatory networks (GRNs), that define neural crest cell identity during different phases of development (Bronner and Simões-Costa, 2016; Martik and Bronner, 2017; Rogers and Nie, 2018), much less is known about how specific GRN components regulate metabolic effector proteins, such as those involved in altering lipid content to control spatiotemporally regulated cellular behaviors. This is a critical question given that metabolic regulation has been shown to be a key effector of neural crest development (Bhattacharya et al., 2021, 2020; Keuls et al., 2020; Nekooie-Marnany et al., 2022; Pascual et al., 2023; Piacentino et al., 2022)

To examine this question, we have focused on the *SMPD3* gene to identify three *cis*-regulatory regions that recapitulate its spatiotemporal expression in the early chicken embryo. Each enhancer regulates expression in different cell types (neural tube versus neural crest) or at different axial levels (cranial versus vagal and trunk). Dissecting the transcriptional inputs into the migratory neural crest-specific *SMPD3* enhancer 3, we show that Sry-box family E (SoxE) transcription factors Sox9 and Sox10 act as upstream regulatory inputs. Finally, we validate that Sox10 regulates endogenous *SMPD3* expression in migratory neural crest cells. These results reveal how developmental GRN components feed into the expression of lipid-metabolizing effector genes during embryogenesis.

## Results

### Identification of putative SMPD3 enhancer sequences

We previously identified the enzyme nSMase2, which hydrolyzes plasma membrane sphingomyelin into ceramide (Shamseddine et al., 2015), as a key regulator of avian neural crest epithelial-to-mesenchymal transition (EMT) where it drives mesenchymalization (Piacentino et al., 2022). Despite the diverse roles and abundance of sphingomyelins and ceramides in all cells (Hannun and Obeid, 2008, 2018), we found that the *SMPD3*, the gene encoding nSMase2, is expressed in a specific spatiotemporal pattern during neurulation that begins at the time of neural crest cell specification (Hamburger Hamilton stage 8, HH8). *SMPD3* transcripts were detected by hybridization chain reaction (HCR) in three major cellular populations along the entire length of the anterior-posterior axis – the neural crest, the neural tube, and the notochord (Piacentino et al., 2022) (**Fig. 1A**).

**Fig 1.**
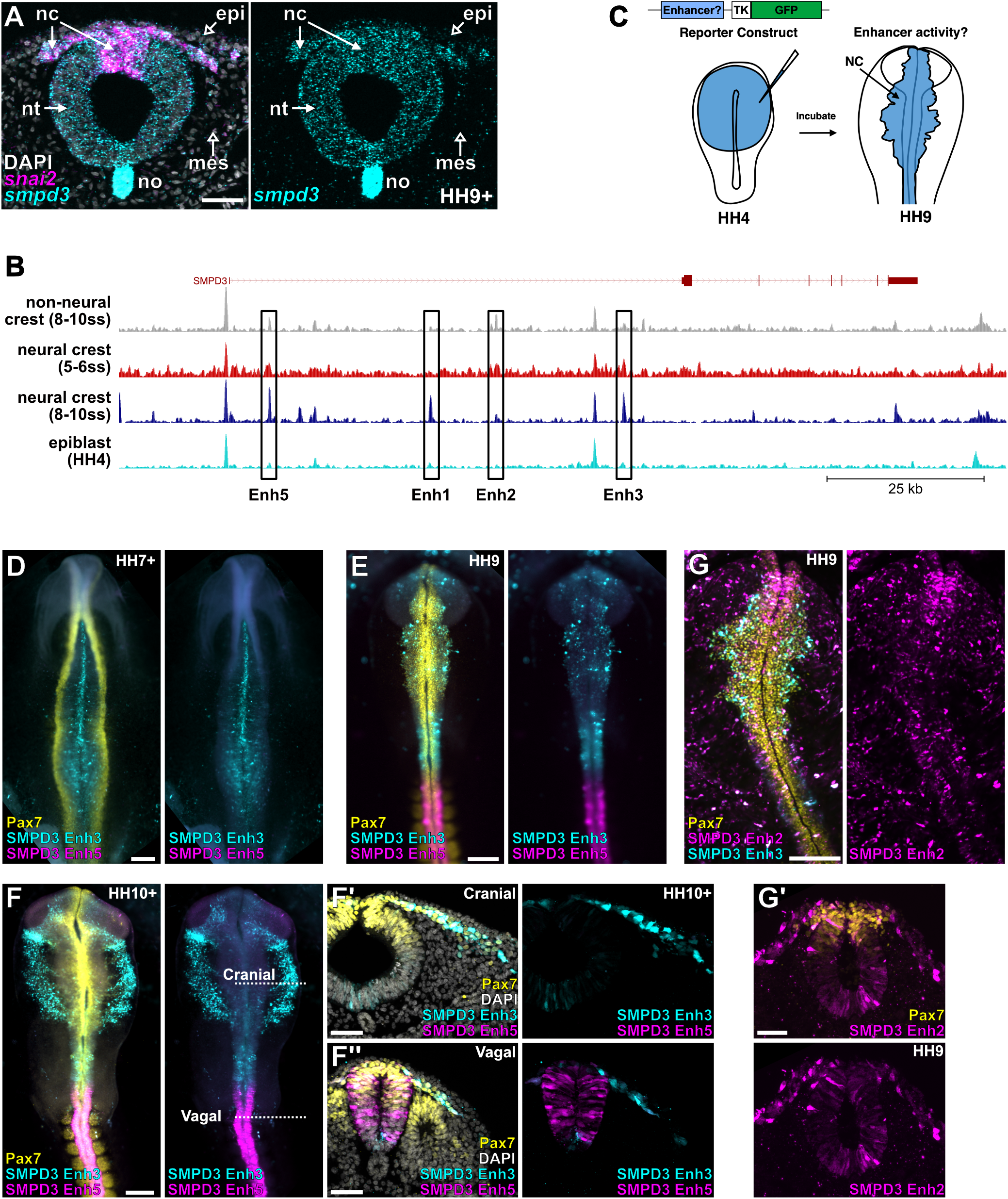
Identification of active regulatory regions in the SMPD3 locus. **A.** Hybridization chain reaction (HCR) reveals the spatial expression pattern of *smpd3* (cyan) and *snai2* (magenta) mRNAs in the chick embryo during early neural crest migration. *Smpd3* expression is present in the developing neural tube (nt), notochord (no), and neural crest (nc), while largely absent from the head mesenchyme (mes) and overlying epidermis (epi). **B.** ATAC-Seq data around the *SMPD3* genomic locus reveal four putative enhancer sequences (boxed) enriched in neural crest over the non-neural crest or epiblast samples. Raw data are drawn from (Williams et al., 2019). **C.** Schematic representing enhancer testing pipeline. Putative enhancers were ligated upstream of the TK basal promoter and assayed for the ability to express a fluorescent protein. These reporter constructs were transfected into the entire epiblast and assayed for fluorescent protein expression at various stages of development. **D-F.** Immunohistochemistry of embryos transfected with SMPD3 Enh3::GFP (cyan) and Enh5::RFP (magenta) at the indicated stages and co-labeled for Pax7 (yellow). F’ and F’’ represent transverse sections of the embryo in F at the indicated axial levels. **G.** Immunohistochemistry for Pax7 in embryos transfected with SMPD3 Enh2::GFP (magenta) and Enh3:: RFP (cyan). G’ represents transverse sections of the embryo in G at the indicated axial level. Scale bars represent 50 µm (A, F’, F’’, G’) and 200 µm (D, E, F, G).

To better understand how sphingomyelin hydrolysis is spatiotemporally regulated, we set out to identify and dissect *cis*-regulatory elements associated with regulating *SMPD3* expression. We began by investigating global changes in accessible chromatin during early neural crest development as profiled by ATAC-seq (Williams et al., 2019). We focused our analysis on peaks of open chromatin that are in close proximity to the *SMPD3* genomic locus and are enriched within neural crest populations over non-neural crest cells. By comparing accessible chromatin from fluorescently-sorted premigratory (Hamburger Hamilton stage (HH) 8+ to HH9) and migratory neural crest cells (HH9+ to HH10), non-neural crest cells (HH9+ to HH10), and epiblast cells during earlier gastrulation (HH4), we identified four non-coding regions with intriguing accessibility across these datasets, each located within the first intronic region of *SMPD3* (**Fig. 1B**). We pursued these sequences since they showed enrichment in neural crest cell populations over epiblast and non-neural crest cells (putative enhancers 1, 3, and 5), or showed accessibility in neural crest and non-neural crest but not epiblast (putative enhancer 2).

### SMPD3 expression is regulated by at least three enhancers during neurulation

We next took advantage of the avian electroporation system which allows for rapid cloning and transfection of putative enhancer sequences to determine regulatory activity (Betancur et al., 2010; Simões-Costa et al., 2012). We cloned these four putative enhancer sequences upstream of a minimal promoter (TK) sequence to drive expression of a fluorescent protein (**Fig. 1C**). Embryos were incubated to select stages ranging from neural crest specification through delamination and migration (HH8 through HH13) and screened for fluorescent reporter expression. From these experiments, we observed fluorescent signal driven by Enhancers 2, 3, and 5. Remarkably, each enhancer construct displayed activity in distinct spatiotemporal patterns (**Fig. 1D-G**), suggesting they work cooperatively to recapitulate the endogenous expression profile of *SMPD3*.

*SMPD3* Enhancer 3 (Enh3) mediated expression first within the developing neural plate, enriched at the midline corresponding to the presumptive floor plate of the neural tube (**Fig. 1D**). While this enhancer showed no activity in the neural crest during specification, it became active in cranial neural crest cells of the midbrain level during early phases of migration at HH9, as well as in the hindbrain neural tube (**Fig. 1E**). During later stages of neural crest migration (HH13), Enh3 exhibited expression broadly in the migratory neural crest cells in the anterior (cranial) and more posterior (vagal) axial levels (**Fig. 1F**). Conversely, Enh5 showed activity in the neural tube starting at the level of the first somite and extending posteriorly at HH9 (**Fig. 1E**). At later stages, Enh5 activity was retained at vagal and trunk axial levels, and persisted in the early migrating vagal neural crest, co-expressed with Enh3 at this level (**Fig. 1F**). Transverse sectioning revealed that Enh5 was active in the premigratory and migratory neural crest, but also in the vagal neural tube (**Fig. 1F’**). Finally, Enh2 showed more unrestricted activity in ectodermal derivatives, including the neural tube and premigratory neural crest at all axial levels, but also in the overlying epidermis (**Fig. 1G**). Together these experiments reveal three enhancer sequences that recapitulate the spatiotemporal expression profile of *SMPD3* in the developing neural tube and neural crest: Enh2 and Enh5 regulate expression in the neural tube and premigratory crest in the anterior and posterior regions, respectively, while Enh3 regulates expression in the migratory neural crest along the entire anterior-posterior axis (**Fig. 1D-G**).

### SoxE factors regulate the expression of SMPD3 Enh3 during neural crest migration

We previously found that *SMPD3* expression is initiated in the neural crest during specification to enable endocytosis and cell signaling necessary for EMT and delamination; however, to our surprise, *SMPD3* transcripts were sustained at high levels during neural crest migration after EMT is complete (**Fig. 1A**) (Piacentino et al., 2022). The activity of Enh3 — which drives reporter expression specifically in migrating neural crest cells (**Fig. 1E-1F**) — suggests that maintained *SMPD3* expression is not simply a consequence of transcript perdurance after an earlier phase of expression. Instead, its activity reveals a distinct, later expression profile and suggests that active *SMPD3* expression during migration may play a subsequent role in neural crest development.

We were intrigued by this finding and sought to understand the upstream regulatory control that feeds into Enh3 activity. Given the timing of Enh3 activity, we hypothesized it may be regulated by members of the SRY-related HMG-box family E (SoxE), such as Sox9 or Sox10, which play important roles in neural crest specification and migration, respectively (Martik and Bronner, 2017). Since *SOX9* expression precedes that of *SOX10,* and *SOX10* expression starts shortly before SMPD3 Enh3 activation, we examined co-expression of *SOX10, SMPD3*, and SMPD3 Enh3 (**Fig. 2A-B**). We confirmed by HCR analysis that *SMPD3* and *SOX10* are co-expressed in the migrating neural crest cells during neural crest migration (**Fig. 2A**), and that SMPD3 Enh3 is active in this *SMPD3* and *SOX10* double-positive population (**Fig. 2B**). These results position Sox10 as a potential regulatory input into SMPD3 Enh3.

**Fig 2.**
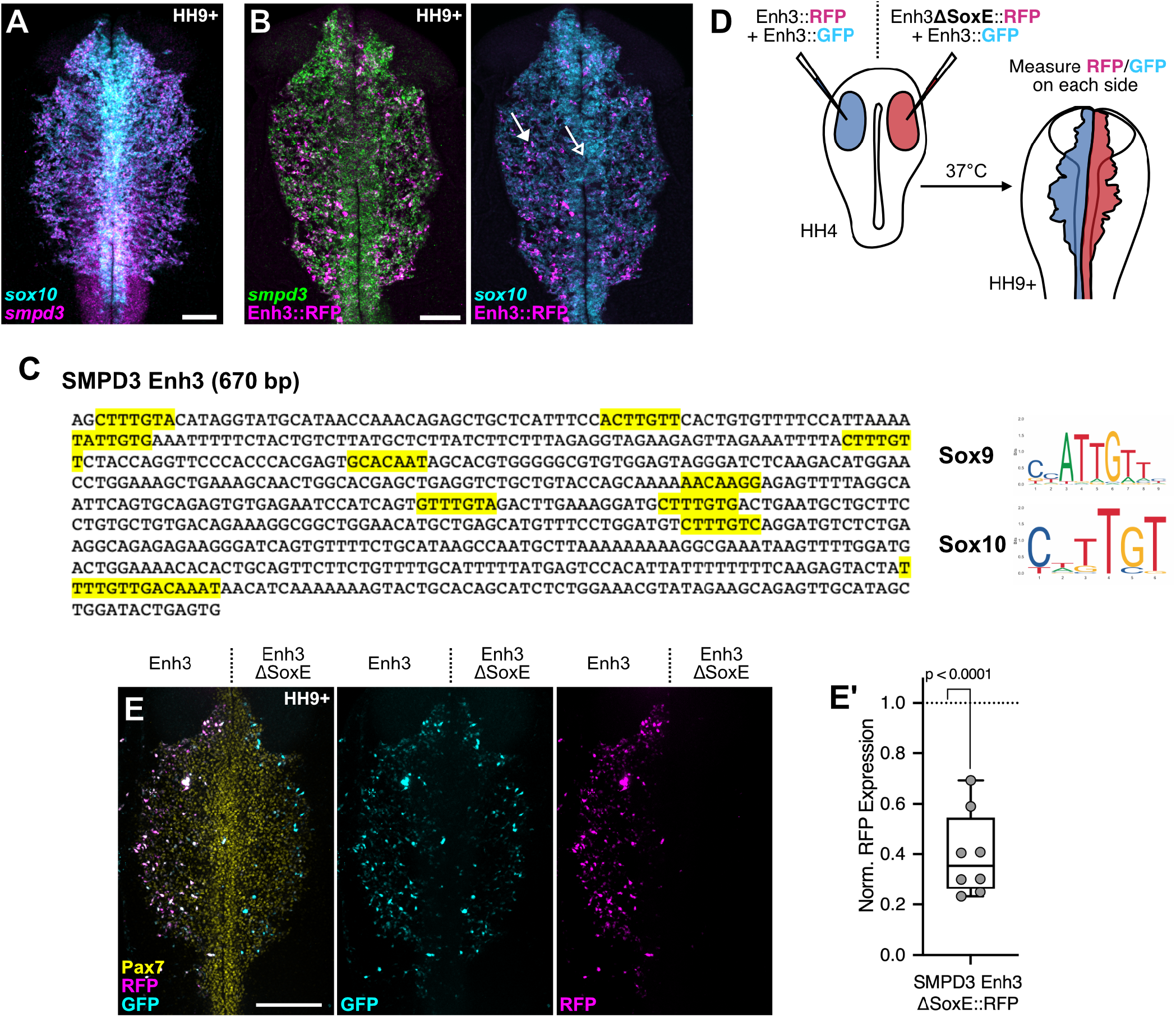
SoxE binding sites regulate the activity of SMPD3 Enh3. **A.** HCR analysis reveals co-expression of the SoxE transcription factor *sox10* (cyan) with *smpd3* (magenta) in the migrating cranial neural crest. **B.** SMPD3 Enh3 (magenta) is expressed in a subset of *smpd3* (green)- and *sox10* (cyan)-expressing neural crest cells. Notably, Enh3::RFP is detected in migrating *sox10+* cells (closed arrow), while it is absent from the more midline *sox10+* premigratory neural crest (open arrow) **C.** SMPD3 Enh3 sequence with putative SoxE transcription factor binding motifs highlighted yellow. Displayed are the Sox9 and Sox10 consensus binding sequences (JASPAR Database (Castro-Mondragon et al., 2022)). **D.** Schematic representing the experimental design to test the role of the SoxE binding sites in SMPD3 Enh3. Both sides were transfected with the wild-type Enh3::GFP. The left side (control) was also transfected with the wild-type Enh3::RFP, while the right side (experimental) was transfected with the Enh3 with the mutated SoxE binding sites (Enh3 ΔSoxE). **E.** Immunohistochemistry reveals the activity of Enh3 and Enh3 ΔSoxE in migratory neural crest (Pax7, yellow) at HH9+. **E’.** Box and whisker quantitation of relative RFP expression normalized to Enh3::GFP expression as a transfection control. Presented are ratios of normalized Enh3 ΔSoxE divided by wild-type Enh3 activity. Each point represents an individual embryo (n=8), and the *p*-value was determined by a two-tailed, paired t-test. Scale bars represent 200 µm.

We next scanned the 670 bp Enh3 sequence for putative transcription factor (TF) binding sites and prioritized TFs with conserved roles in the neural crest gene regulatory network (GRN)(Martik and Bronner, 2017). By examining predicted TF binding site motif enrichment as determined by Williams and colleagues (**Fig. S1**)(Williams et al., 2019), we found four larger predicted SoxE family binding motifs (**Fig. S1**), and a total of 11 sites that fit the core Sox9 and Sox10 signature (**Fig 2C**).

The roles of SoxE-family TFs are highly conserved during neural crest specification and migration, so we asked if their input into Enh3 is required for its activity. To eliminate SoxE binding to Enh3, we mutated the 11 SoxE binding signatures and produced an Enh3ΔSoxE::RFP reporter. We transfected this mutant construct on the right side and wild type Enh3::RFP on the left side of chicken embryos (**Fig. 2D**). We included wild type Enh3::GFP on both sides to normalize for electroporation efficiency. Enh3ΔSoxE::RFP showed negligible expression compared to the wild-type enhancer (**Fig. 2E**), indicating that these SoxE binding sites are necessary for Enh3 activity.

### SOX10 is a primary regulator of SMPD3 Enh3 during neural crest migration

The SoxE family consists of three TFs — Sox8, Sox9, and Sox10 — and we next asked which of these TFs regulates Enh3 activity. Sox8 is expressed first during specification at early HH8 stages (Betancur et al., 2011), followed by Sox9 during late HH8, and finally Sox10 during HH9 and the onset of delamination and migration (Betancur et al., 2011, 2010; Cheung and Briscoe, 2003; McKeown et al., 2005). Enh3 activity was not detected in the midline where *SOX10* expression begins just prior to delamination (**Fig. 2B**, open arrow), but is then active after delamination in the more lateral, migrating *SOX10*+ neural crest cells (**Fig. 3A**, filled arrow). Given this expression pattern, we predicted that Sox10, and not earlier Sox9, is the most likely candidate that regulates Enh3 activity. To test this prediction, we knocked out Sox10 using CRISPR/Cas9 (Gandhi et al., 2021), and compared Enh3::RFP activity on control and Sox10-knockout sides. We found that Sox10 knockout was sufficient to significantly diminish activity from Enh3, indicating that Sox10 indeed acts upstream of SMPD3 Enh3 (**Fig. 3A**). Similarly, we found that Sox9 knockdown by translation blocking antisense morpholino also diminished activity from Enh3 (**Fig. 3B**). We also observed reduced Sox10 protein levels following Sox9 knockdown, consistent with previous reports on the role of Sox9 in regulating *SOX10* expression (Betancur et al., 2010). Taken together, our results suggest that Sox10 is directly upstream *SMPD3*, while Sox9 regulates *SMPD3* indirectly through *SOX10*.

**Fig 3.**
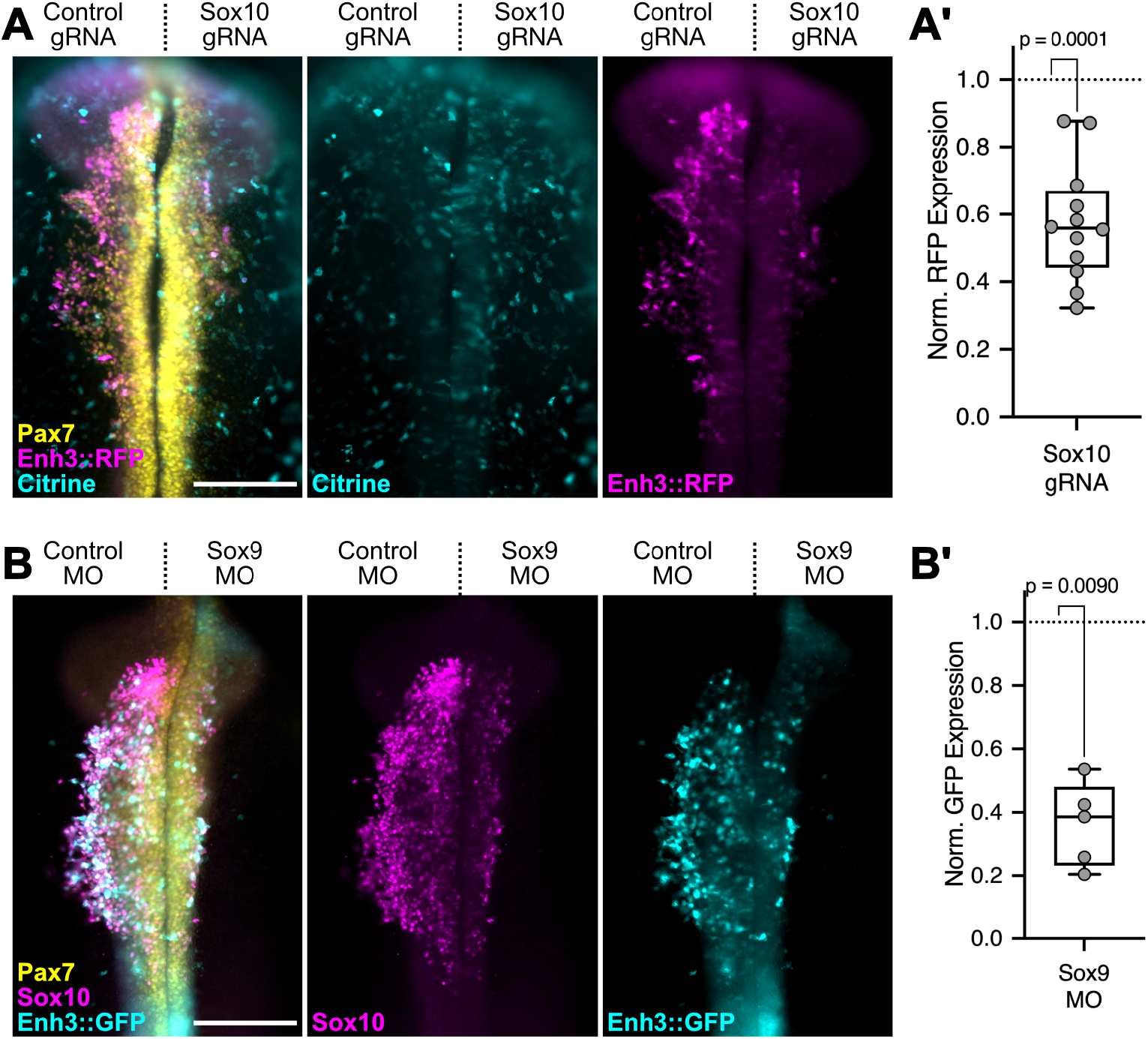
Sox10 and Sox9 are upstream of SMPD3 Enh3 activity. **A.** Embryos were transfected with SMPD3 Enh3:: RFP (magenta) together with a control (left) or Sox10-targeted (right) CRISPR construct (expressing Cas9, gRNA, and Citrine fluorescent protein). The fluorescent intensity of RFP was measured on both sides and normalized to Citrine as a transfection control. The results are displayed as a box and whisker plot revealing the relative RFP expression following Sox10 knockdown divided by control (A’). **B.** SMPD3 Enh3::GFP activity was assayed after co-transfection with control and Sox9-targeting morpholinos. The fluorescent intensity of RFP was measured on both sides and normalized to Citrine as a transfection control. The results are displayed as a box and whisker plot revealing the relative GFP expression following Sox9 knockdown divided by control (B’). Each point represents an individual embryo (n=12 Sox10 gRNA; n=5 Sox9 MO), and *p-*values were determined by two-tailed, paired t-tests. Scale bars represent 200 µm.

### SOX10 regulates endogenous SMPD3 expression during neural crest migration

Finally, we sought to determine if Sox10-dependent Enh3 activity reflects endogenous *SMPD3* expression in the migratory neural crest. Given that endogenous *SMPD3* expression precedes that of *SOX10*, we expected earlier *smpd3* transcripts to perdure through early neural crest migratory stages (HH9-HH10), masking any Sox10-dependent changes at this stage *in vivo*. To overcome this limitation, we performed Sox10 knockdown using a translation-blocking morpholino introduced at HH4, explanted neural crest cells prior to delamination at HH8+, and cultured explanted neural crest cells for 16 hours to allow for cell dispersion as a monolayer of migratory neural crest cells, providing ample time for “premigratory” *smpd3* expression to diminish and “migratory” *smpd3* expression to take over. We performed HCR for *smpd3* on the explanted neural crest cells and measured transcript abundance. The results show a significant reduction in *smpd3* expression following Sox10 knockdown (**Fig. 3A-C**), supporting the conclusion that Sox10 regulates *SMPD3* expression during neural crest migration.

## Discussion

Here we have identified three enhancers that recapitulate the spatiotemporal expression profile of *SMPD3* in the ectodermal lineages during avian embryo neurulation. *SMPD3* Enh3 drives expression specifically in the migrating neural crest, while Enh2 and Enh5 regulate the *SMPD3* expression observed in the developing neural tube (**Fig. 1**). By examining predicted transcription factor binding sites across these three *SMPD3* enhancers (**Fig. S1–S3**), we present a model of validated and potential transcriptional regulators that determine the precise *SMPD3* expression domains (**Fig. 4E**). By generating binding site mutations in the Enh3 reporter construct (**Fig. 2**) and knocking down upstream transcription factors (**Fig. 3**), we determined that Sox9 and Sox10 act to positively regulate the activity of *SMPD3* Enh3 during neural crest migration. Given the role of Sox9 in regulating the expression of Sox10 (Betancur et al., 2010), and the onset of *SMPD3* Enh3 activity shortly after *SOX10* expression (**Fig. 2B**), we hypothesize that Sox10 regulates Enh3 through a direct interaction, while Sox9 acts indirectly and upstream of Sox10. Finally, we show that Sox10 knockdown reduces the expression of endogenous *SMPD3* in late-migrating neural crest cells (**Fig. 4A-4C**), validating the relevance of this transcriptional interaction. Together our results reveal a novel mode of transcriptional regulation of *SMPD3* gene expression.

**Fig 4.**
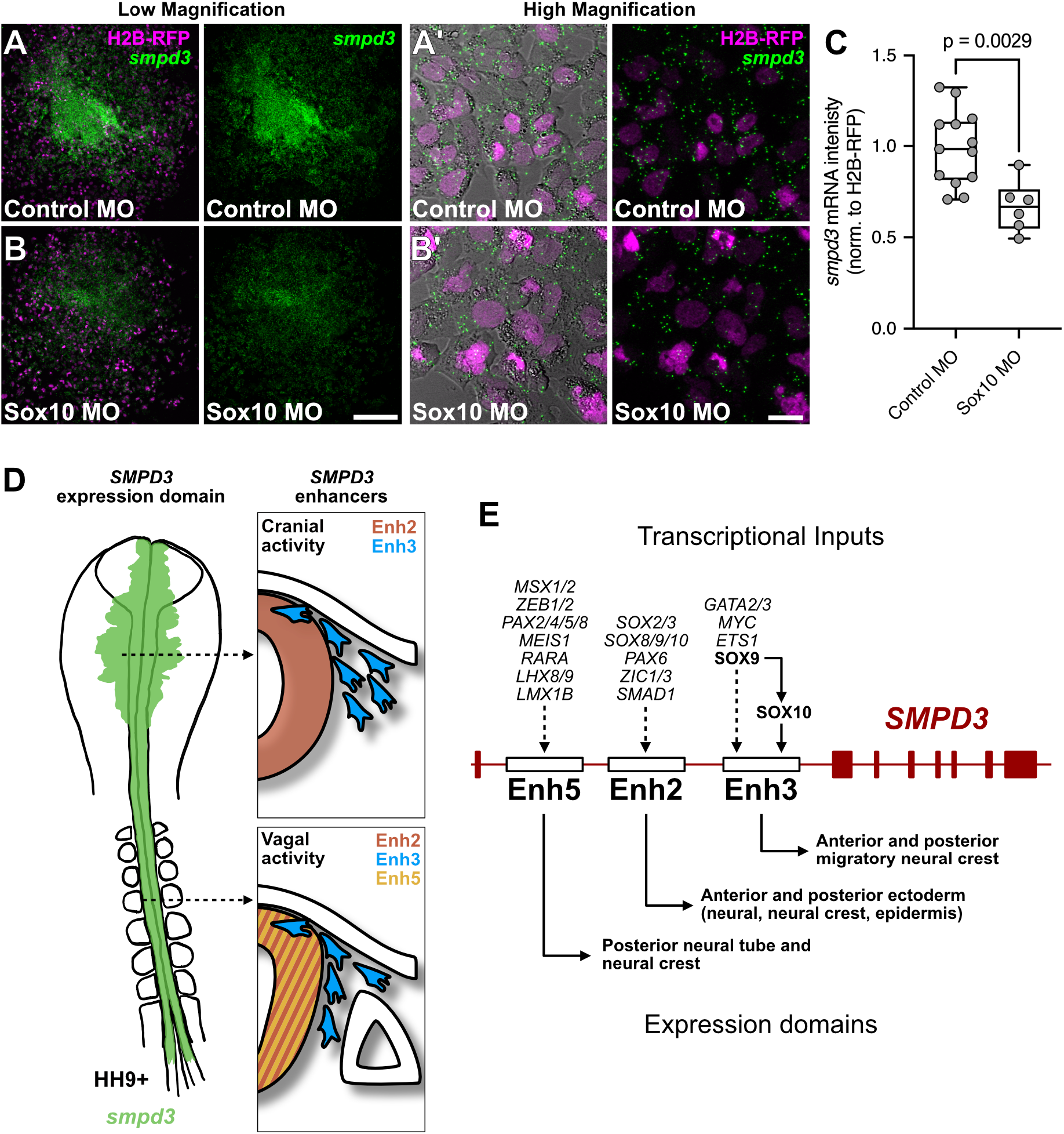
Sox10 knockdown provokes late loss of *smpd3* expression. Embryos were transfected during gastrulation with an H2B-RFP-expressing construct (magenta), and neural crest cells were explanted into a two-dimensional culture system during neurulation. Explants that received non-targeted control (A) and Sox10-targeting morpholinos (B) were allowed to adhere and migrate overnight at which point they were subjected to HCR to detect endogenous *smpd3* transcripts. Displayed are low magnification views (A, B) showing the full explants and high magnification views (A’, B’) showing individual cells. **C.** *Smpd3* fluorescence intensity was quantitated and normalized to H2B-RFP for Control explants and Sox10 MO explants to reveal a significant reduction in *smpd3* expression following Sox10 knockdown. Each point represents an individual explant (n=13 Control MO; n=6 Sox10 MO), and a two-tailed, unpaired t-test determined the *p*-value. Scale bars represent 200 µm (A, B) and 20 µm (A’, B’). **D.** Schematic representation of endogenous SMPD3 gene expression (left, green) along the anterior-posterior axis, with Enh2, Enh3, and Enh5 expression domains as indicated at the cranial and vagal levels. **E.** Map of the SMPD3 genomic locus displaying the relative positions of Enhancers 2, 3, and 5 in the first intron and summarizing their upstream transcriptional inputs and downstream domains of expression. Bold transcription factor names are functionally tested herein, while italicized names are predicted from binding site analyses.

In addition to the migratory neural crest-specific *SMPD3* Enh3, we found that *SMPD3* expression in the neural tube was controlled by two separate enhancers, Enh2 and Enh5 (**Fig. 1**). This neural tube expression is subdivided axially with Enh2 driving expression along the full anterior-posterior axis, and Enh5 limited to just the posterior neural tube corresponding to the presumptive spinal cord. These expression domains are consistent with many of their predicted transcriptional input sites (**Fig. S2–S3**)(Williams et al., 2019). Notably, posterior expression of Enh5 is likely regulated by transcription factors enriched in the developing posterior neural tube including *MEIS1/2* (Sánchez-Guardado et al., 2011), *MSX1/2* (Anderson et al., 2019; Khudyakov and Bronner-Fraser, 2009), *ZEB1/2* (Dady et al., 2012; Rogers et al., 2013), and by retinoic acid signaling coming from *RALDH2* expression in the adjacent somites (Niederreither and Dollé, 2008; Olivera-Martinez and Storey, 2007). While Enh2, which is expressed both posteriorly and anteriorly, similarly contains a *MEIS1/2* binding site, it also has potential inputs from *SOX2/3* and *ZIC1/3* which are expressed in the full length of the neural tube (Khudyakov and Bronner-Fraser, 2009; Odaka et al., 2018; Rex et al., 1997), and from more anteriorly-restricted *LMX1B* and *PAX6* (Adams et al., 2000; Bhattacharyya et al., 2004; Yuan and Schoenwolf, 1999). Since *SMPD3* expression is highly enriched in the mammalian central nervous system (Hofmann et al., 2000; Stoffel et al., 2018) at adult stages, our results suggest a regulatory scheme that establishes this expression profile during early development.

In addition to the neural tube, we observed SMPD3 Enh2 expression in the presumptive epidermis, which lacks endogenous *SMPD3* expression (compare **Fig. 1A** and **Fig. 1G**). This may be regulated in part by *PAX6* and *SMAD1* inputs in the epidermis (Bhattacharyya et al., 2004; Faure et al., 2002; Piacentino and Bronner, 2018). However, since endogenous *SMPD3* is not expressed in the epidermis, we hypothesize that a separate regulatory element acts to suppress these interactions; without these negative inputs, our reporter construct thus displays an ectopic expression domain. Finally, our electroporation experiments robustly transfect ectodermal tissues but not the prospective notochord; therefore, we cannot conclude if these or other enhancers act to drive the *SMPD3* expression observed in the developing notochord.

Post-translational mechanisms that regulate the localization and activity of the *SMPD3* gene product, nSMase2, have been well-studied (Airola et al., 2017; Shamseddine et al., 2015; Tani and Hannun, 2007; Wu et al., 2010). For example, binding to phosphatidylserine in the plasma membrane triggers a conformational change that locally activates the nSMase2 catalytic domain (Airola et al., 2017). However, much of our knowledge of *SMPD3* transcriptional regulation is determined from experiments in cell lines. For example, in mouse chondrocytes, loss of the SoxE transcription factor, Sox9, diminished *SMPD3* expression levels, and Sox9 was found to directly bind two regions in the first intron of the rat *SMPD3* locus, thereby upregulating activity from the *SMPD3* basal promoter (Li et al., 2016). Our results also show SoxE-mediated regulation through *SMPD3* Enh3, and potentially Enh2 (**Fig. S2**), demonstrating a highly conserved regulatory program likely activated earlier in development, prior to this later role in chondrocyte differentiation. Furthermore, *SMPD3* expression is induced by retinoic acid signaling through *ATRA* in stress-induced growth arrest in breast cancer cell lines (Clarke et al., 2016), and can be induced by *BMP2* signaling through the activation of *RUNX2* in C2C12 myoblasts (Chae et al., 2009). Putative retinoic acid receptor α (*RARA*) and BMP-responsive *SMAD1* binding sites in *SMPD3* Enh5 and Enh2, respectively, are consistent with these inputs also participating in developmental expression. While these studies indicate regulatory interactions that induce *SMPD3* expression in response to external stimuli, our results reveal the regulatory logic underlying endogenous *SMPD3* regulation during normal *in vivo* embryonic development.

Taken together, our results show how developmental gene regulatory networks, such as those present within the developing vertebrate neural crest, regulate specific expression domains of the metabolic lipid effector gene *SMPD3*. We found that sphingolipid metabolism is spatially regulated by two different families of Sox genes, with SoxB1 members like *SOX2* and *SOX3* regulating expression in the neural tube through *SMPD3* Enh2, and SoxE members including *SOX9* and *SOX10* maintaining expression in the neural crest through *SMPD3* Enh3. We posit that determining transcriptional inputs that drive specific expression patterns of key enzymes involved in lipid metabolism will reveal core signals, transcription factors, and chromatin remodelers that work in concert to drive a suite of metabolic regulators, thereby resulting in large-scale changes in the cellular lipidome during developmental transitions such as epithelial-to-mesenchymal transition or differentiation.

## Acknowledgments

We would like to thank Drs. Tatjana Sauka-Spengler, Ruth Williams, and Ivan Candido Ferreira for sharing data and helpful discussion on enhancer identification and dissection. We’d also like to thank the Caltech Biological Imaging Facility, supported by the Caltech Beckman Institute and the Arnold and Mabel Beckman Foundation, for confocal microscopy support. Funding for this work comes from the NIH grants K99/R00 DE029240 to M.L.P., R01 DE027538 to M.E.B., and a travel award provided by the Company of Biologists to M.L.P.

## Author Contributions

Conceptualization: M.L.P.

Experiment design: M.L.P.

Experimentation: M.L.P., A.J.F., A.C.A., I.G., and M.E.B.

Data analysis: M.L.P. and A.J.F.

Data interpretation: M.L.P. and M.E.B.

Manuscript preparation: M.L.P.

Manuscript editing: M.E.B.

## Materials and Methods

### Enhancer identification and cloning

To identify putative enhancer regions around the *SMPD3* locus, we aligned previously processed and published ATAC-Seq reads (Williams et al., 2019) to the chicken genome (galGal4 assembly) using the University of California, Santa Cruz, Genome Browser (Kent et al., 2002), and examined the vicinity around the *SMPD3* gene locus on chromosome 11. We visually compared ATAC-Seq peaks from purified neural crest cells from premigratory stages (5-6 somite stage, ss), migratory stages (8-10ss), non-neural crest cells from the same stages, and unsorted epiblast cells from Hamburger Hamilton stage HH4 (**Fig. 1B**). Identification of putative transcription factor binding sites was performed manually by scanning for the core SoxE binding motif from JASPAR ((Castro-Mondragon et al., 2022), **Fig. 2C**), or by examining previously determined predicted transcription factor binding sites (Williams et al., 2019) and manually annotating using the Benchling sequence analysis toolkit (https://www.benchling.com).

Peaks enriched in neural crest samples were cloned from genomic DNA using standard PCR methods using AccuPrime Hi-Fidelity Taq polymerase (Invitrogen). Primers used to amplify target enhancer sequences are provided in the Key Resources table (**Table S1**). Amplified enhancers were then digested using BsmBI and ligated into a reporter construct contining the basal TK promoter driving expression of Citrine fluorescent protein (Williams et al., 2019). For Enh3 and Enh5 constructs, the Citrine fluorescent protein was replaced with EGFP or mRFP by digestion with NheI and XbaI, following standard subcloning methods. The *SMPD3* Enh3ΔSoxE construct was generated by synthesizing a gBlock gene fragment in which each of 11 putative SoxE binding motifs was replaced with a scrambled sequence, indicated in Table S1, and was subcloned into the mRFP-containing reporter construct as described above. All plasmids produced in this study were verified by Sanger sequencing (Laragen) or full plasmid sequencing (Primordium Lab) prior to use.

### Embryos and perturbations

Fertilized chicken eggs were obtained from commercial sources (Sunstate Ranch, Sylmar, CA and UConn Poultry Farm, Storrs, CT), and incubated at 37.5°C to the desired Hamburger Hamilton (HH) stages (Hamburger and Hamilton, 1951). Electroporations were performed *ex ovo* at gastrula stages (HH4) using five pulses of 5.6 V for 50 ms at 100 ms intervals and cultured to the desired stages in albumin with 1% penicillin/streptomycin at 37.5°C. All plasmid DNAs were electroporated at 2.5 µg/µl working concentrations (reporter and CRISPR/Cas9 constructs (Gandhi et al., 2021)). All morpholino oligonucleotides were synthesized by GeneTools LLC, and were used at the following doses: Sox9 MO at 0.7 mM (Betancur et al., 2010); Sox10 MO at 1.25 mM (Barembaum and Bronner, 2013); Control MO at 0.7 or 1.25 mM. Morpholino and CRISPR/Cas9 target sequences are described in Table S1.

Embryos of the desired stage or neural crest explants were fixed with 4% paraformaldehyde in phosphate buffer for 10 minutes at room temperature before processing for Hybridization Chain Reaction (HCR) or immunohistochemistry (see below for details). Fixed and labeled embryos were processed for transverse cryosectioning by incubation in 5% sucrose for 15 minutes at room temperature, in 15% sucrose overnight at 4°C, then in 7.5% gelatin in 15% sucrose at 37.5°C overnight, before embedding and flash freezing with liquid nitrogen. Samples were then stored at −80°C before cryosectioning at a thickness of 18 µm.

### Hybridization chain reaction

Hybridization chain reaction (HCR) labeling kits were purchased from Molecular Technologies, and all experiments were performed following the manufacturer’s instructions (Choi et al., 2018). Probe sets used in this study include *smpd3* (B1 initiator), *snai2* (B4), *sox10* (B3), and were detected using appropriate amplifier hairpins labeled with Alexa488, Alexa546, or Alexa647.

### Immunohistochemistry

After fixation, embryos were washed into TBST + Ca^2+^ (50 mM Tris-HCl, 150 mM NaCl, 1 mM CaCl_2_, 0.5% Triton X-100), and all blocking and labeling steps were made in this buffer. Blocking was performed in 10% donkey serum for 1 hour at room temperature, and primary and secondary antibody incubations were carried out in 10% donkey serum for 2 nights each at 4°C. Primary antibodies used here include mouse IgG1 anti-Pax7 (1:10; DSHB #PAX7), rabbit polyclonal anti-Sox10 (1:500; Atlas Antibodies #HPA068898), goat polyclonal anti-GFP (1:500; Rockland #600-101-215M), and rabbit polyclonal anti-RFP (1:500; MBL #PM005). Primary antibodies were then counter-labeled with Alexa Fluor 350-, 488-, 568-, 633-, or 647-conjugated donkey secondary antibodies (1:500; Molecular Probes).

### Explant culture

Neural crest explants were prepared from control and Sox10 MO-electroporated embryos as previously described (Piacentino et al., 2021). Polymer coverslip imaging chambers (ibidi) were coated with 25 µg/mL fibronectin in PBS for 1 hour at 37°C, then aspirated and replaced with explant culture media (L15 supplemented with 10% Fetal Bovine Serum, 10% Chick Embryo Extract, and 1% Penicillin/Streptomycin). Control and Sox10-depleted dorsal neural tubes were manually dissected from the midbrain level of stage HH8+/HH9- embryos using fine scissors, transferred to the imaging chambers, and cultured overnight at 37.5°C before fixation and HCR processing (see above).

### Imaging, analysis and statistics

Whole mount and transverse section imaging was performed using a Zeiss Imager.M2 with an ApoTome.2 module, with a Zeiss LSM 880 upright confocal microscope, or a Nikon AX inverted confocal microscope. Fluorescent whole mount embryo images were collected as wide-field views (**Fig. 1D-F** and **Fig. 3**), or displayed as a maximum intensity projection of Z-stacks (**Fig. 1G**, **Fig. 2A-B**, and **Fig. 2E**). Fluorescent transverse section images are displayed as maximum intensity projections of Z-stacks (**Fig. 1A**, **Fig. 1F’-1F’’**, and **Fig. 1G’**). Explant experiments were imaged using a LSM 980 inverted confocal microscope with a 40x/1.20 NA objective and images are displayed as maximum intensity projections of Z-stacks (**Fig. 4A,B**).

All images were prepared for display and analyzed using Fiji (Schindelin et al., 2012), and fluorescent intensity measurements were made as previously described (Piacentino et al., 2022). All source data and code used for measurements and analysis can be found on GitHub (https://github.com/PiacentinoLab/2023_SMPD3_Transcriptional_Regulation). Briefly, for normalized enhancer activity measurements presented in Figures 2 and 3, regions of interest were drawn around to encompass the neural crest cell area on the left control side and right experimental side of the midbrain. These areas were used to measure fluorescence integrated density for the reporter of interest and a normalization control (wild-type reporter or ubiquitous expression of H2B-RFP). Measured intensities were then normalized to the electroporation control, and then ratios of experimental/control values were determined using R and plotted using GraphPad Prism 9. For neural crest explants in Figure 4, three 175 µm x 175 µm square regions of interest were drawn randomly over the migratory neural crest cell area and used to measure *smpd3* HCR and H2B-RFP electroporation tracer intensity. *smpd3* intensity was normalized against H2B-RFP to control for differences in cell density between regions of interest before the three regions were averaged together to produce one representative value per explant.

All statistical computations were performed using GraphPad Prism 9. All datasets were tested for normal distribution with a Kolmogorov-Smirnov test. Normalized control and experimental intensities from *in vivo* reporter analyses (**Fig. 2** and **Fig. 3**) were analyzed with two-tailed paired *t*-tests. Normalized *smpd3* expression in neural crest explants (**Fig. 4**) were analyzed with a two-tailed unpaired *t*-test. Specific tests and their *p* values are displayed in the relevant figures and legends. Box-and-whisker plots display the ratio of experimental divided by corresponding control measurements (**Fig. 2** and **Fig. 3**), or normalized *smpd3* expression (**Fig. 4**) with each biological replicate displayed as individual points. The box plot elements displayed include: center line, median; box limits, upper and lower quartiles; whiskers, minimum and maximum values.

---

**Fig S1.**
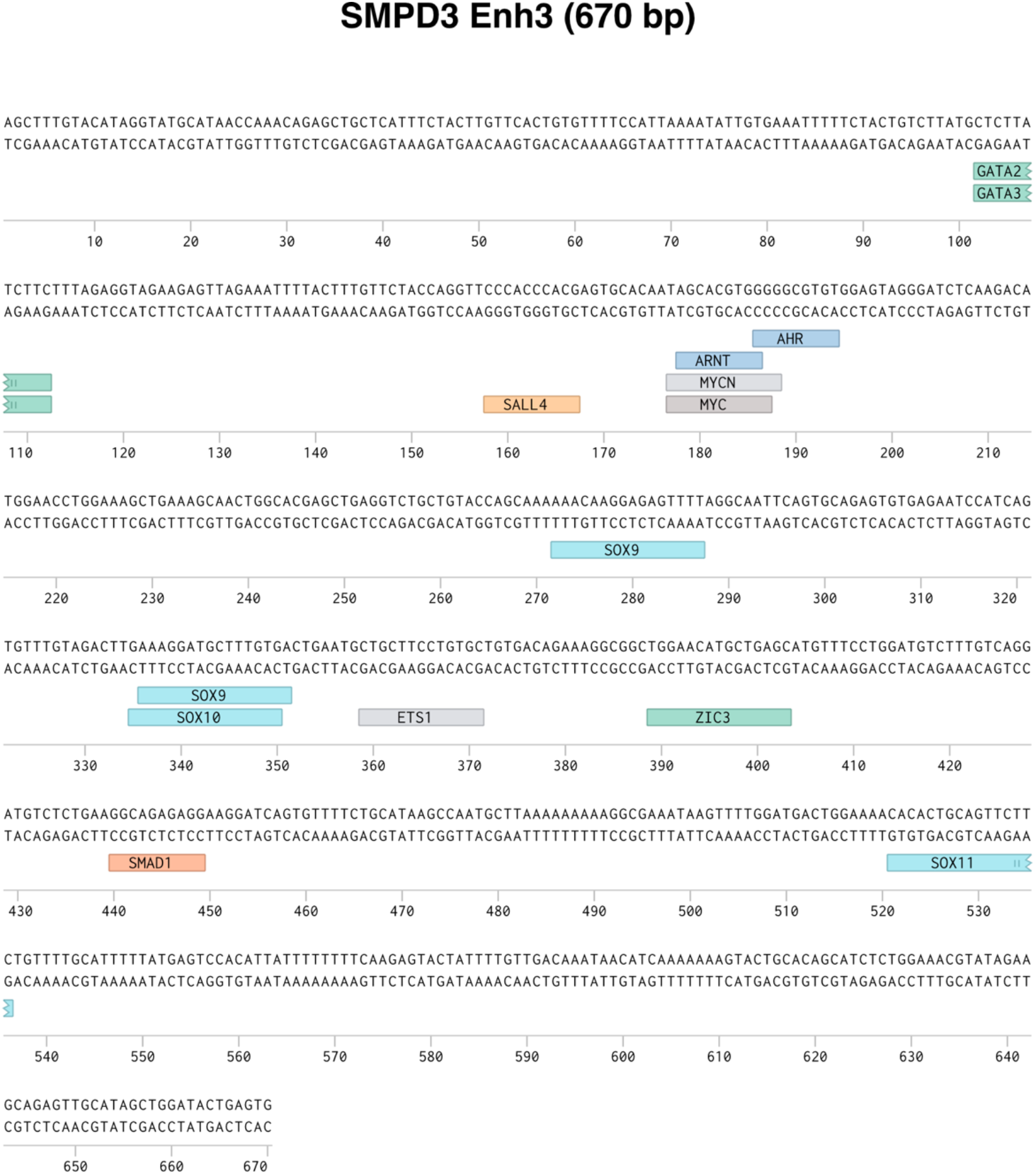
Binding site predictions for migratory neural crest-specific *SMPD3* Enh3. **A.** Transcription factor binding site predictions for the SMPD3 enhancers 3, as drawn from HOMER analysis (Williams et al., 2019).

**Fig S2.**
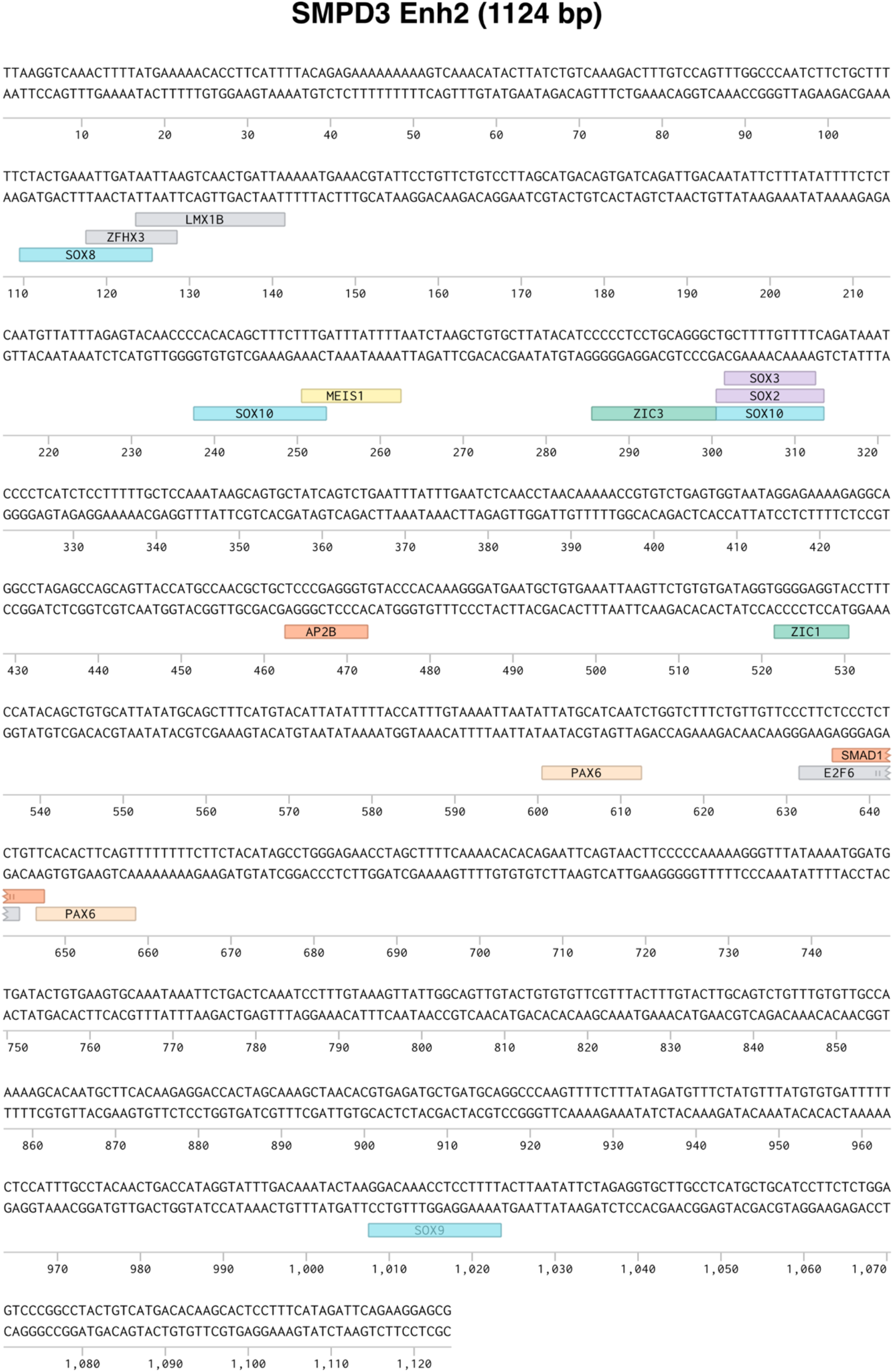
Binding site predictions for ectodermal *SMDP3* Enh2. **A.** Transcription factor binding site predictions for the SMPD3 enhancers 2, as drawn from HOMER analysis (Williams et al., 2019).

**Fig S3.**
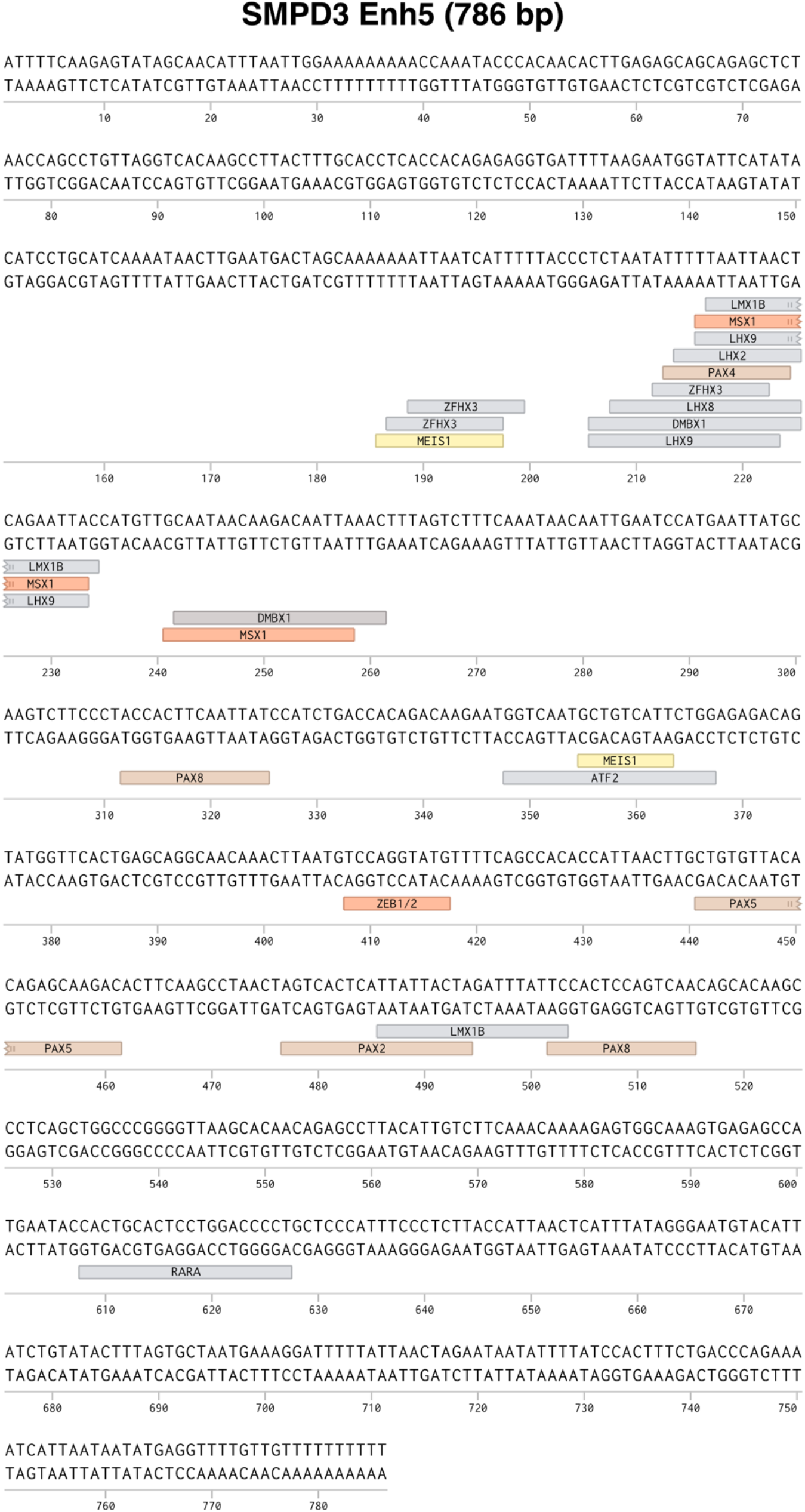
Binding site predictions for posterior neural tube-specific SMPD3 Enh5. **A.** Transcription factor binding site predictions for the SMPD3 enhancers 5, as drawn from HOMER analysis (Williams et al., 2019).

**Table S1.**
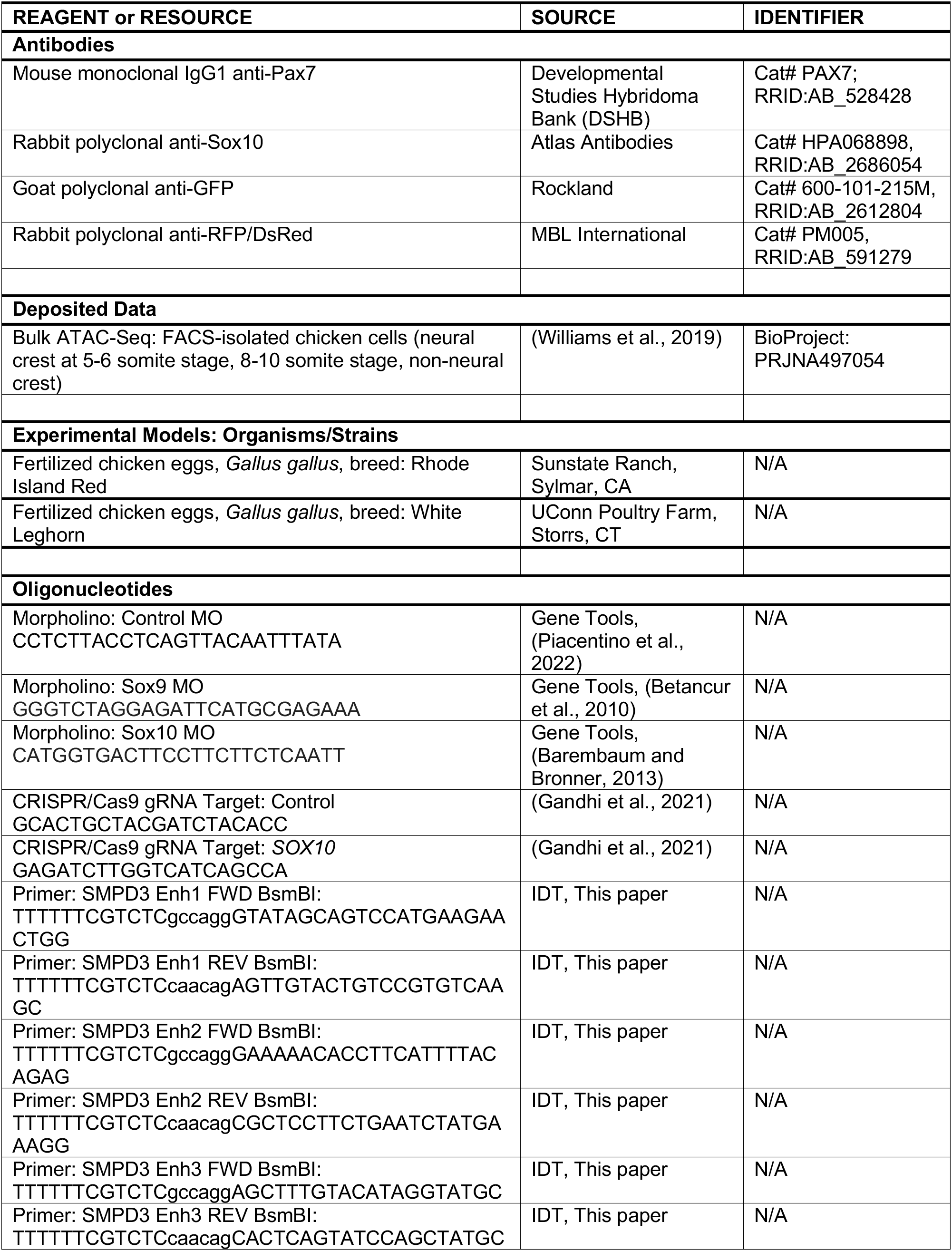
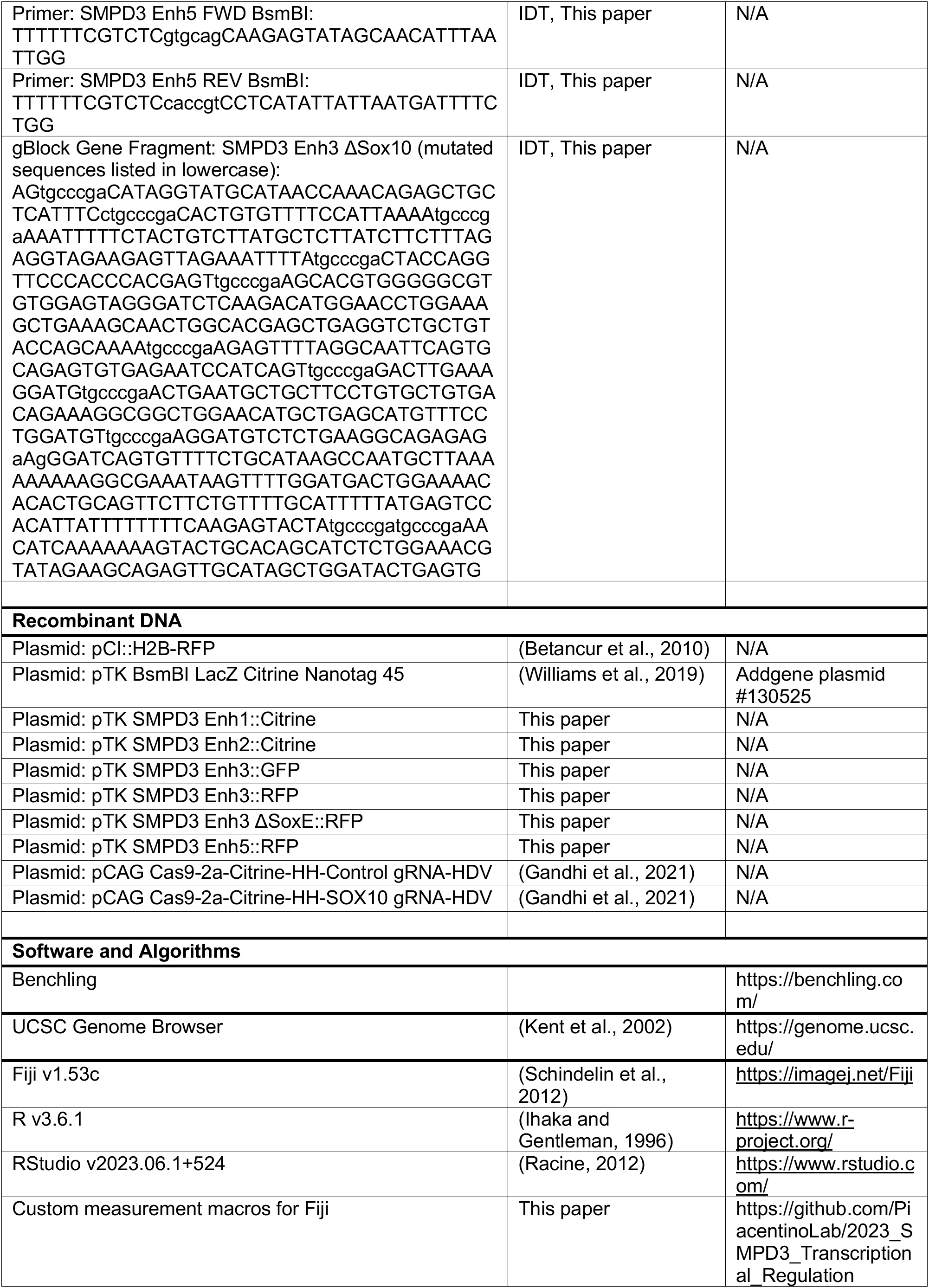
Key Resources.

